# Regeneration leads to global tissue rejuvenation in aging sexual planarians

**DOI:** 10.1101/2023.06.24.546358

**Authors:** Xiaoting Dai, Xinghua Li, Scott Pletcher, David Paris, Leonid Kruglyak, Jacob Sobota, Longhua Guo

## Abstract

An ability to delay aging—or to reverse the negative effects of aging—could prevent age-related disease and greatly enhance quality of life in old age. However, whether it is possible to globally reverse the physiological effects of aging in order to extend healthspan is unknown. The freshwater planarian *Schmidtea mediterranea* has been considered “immortal” due to its exceptional tissue regeneration capabilities. Here, we report that a sexually reproducing lineage of *S. mediterranea* exhibits age-associated physiological decline 12 months after birth. Age-associated changes include alterations in sensory organs, loss of neurons and muscle, loss of fertility, and impaired motility, but no reduction in stem cells at the age of 3 years. Differential gene expression analysis, comparing young and old planarian cells, furthermore revealed cell-type-specific changes in transcription as well as changes in classical aging pathways (e.g., insulin signaling). Remarkably, amputation followed by regeneration of lost tissues led to a global reversal of these age-associated changes. Older individuals that underwent regeneration showed restored youthful patterns of gene expression, stem cell states, tissue composition and rejuvenation of whole-animal physiology. Our work reveals a naturally evolved solution to age reversal in planaria that may provide insights into anti-aging strategies in humans.

## Introduction

Adult stem cells (ASCs) undergo self-renewal and differentiation to replace cells lost to wear and tear. In old age, these regenerative capacities of ASCs diminish^1–5^, and this phenomenon is one of the major causes of frailty and disease^1^. Humans begin losing skeletal muscle mass in their 40s; by the age of 80, up to 50% of muscle mass may be lost, contributing to impaired function and disability^6,7^. Similarly, cell type specific neuronal loss occurs in the human hippocampus, already beginning at the age of 13 and continuing into the 80s^8^. Age-associated loss of neurons is even more devastating in the context of neurodegenerative diseases, such as Alzheimer’s disease^9,10^. A method for rejuvenating the regenerative functions of aged ASCs would revolutionize medicine and significantly aid efforts to combat aging and age-related disease.

Several interventions have been shown to rejuvenate old cells in multiple tissues^2,4^. These include heterochronic parabiosis^11^, exercise^12–14^, caloric restriction^15^, and reprogramming^2,4,16^. However, the aged hematopoietic stem cell system, which is critical to mammalian healthspan and lifespan, remains refractory to these interventions^2,17^ and reprogramming reagents have to be delivered in a tissue-specific fashion^16^. Hence, while rejuvenation research has shown great promise in restoring stem cell or niche functions in multiple tissues, global rejuvenation of all tissues or age-reversal of whole-animal physiology remains elusive.

Long-lived species provide unique opportunities to uncover naturally-evolved mechanisms for the extension of health span and lifespan. For example, studies in the naked mole rat, a long-lived rodent, reported a large quiescent hematopoietic stem cell pool with prolonged cell cycles^18^ relative to the short-lived house mouse, motivating extensive research on stem cell regulation in long-lived species and underscoring its potential to extend lifespan and healthspan.

Planarians constitute a popular long-lived model system. Freshwater planarians have been called “immortal” due to their extremely long lifespan^19–23^ and unique tissue regeneration capabilities ^24,25^. Regeneration is tightly regulated by division and differentiation of abundant adult stem cells (neoblasts), and positional guiding signals expressed by muscle cells^26–28^. In the past two decades, genomic^29,30^ and molecular analyses^31–36^ have provided a mechanistic understanding of regeneration in several planarian species, including *Schmidtea mediterranea*. It was reported that telomeres shorten^37^, and viable progeny decline^38^ in older planarians. Whether planarians experience aging and show a typical age-dependent decline in physiological, cellular and molecular functions has not been systematically examined, in part because of the challenges inherent in measuring lifespan in a long-lived animal, or even defining age in asexual planarians that undergo a vegetative mode of reproduction (i.e., fission^39^). Inbred lines of a sexual lineage of *S. mediterranea* have been established to study genetic variations and chromosome biology^30,40^. This resource provides a unique opportunity to examine aging in this long-lived model system and disentangle genetic control from environmental effects.

Here, we examine aging in the sexual lineage of *S. mediterranea.* To employ this model for aging research, we define chronological age as time since fertilization, thus overcoming the challenges involved in lineages that rely on vegetative reproduction. As in the more traditional short-lived model systems (mice, fish, nematodes and fruit flies), *S. mediterranea* exhibits signs of aging at multiple levels—molecular, cellular and physiological. However, unlike other model systems, we show here that *S. mediterranea* is capable of reversing age-related changes. We show that amputation and regeneration trigger global rejuvenation that completely reverses the multi-level effects of aging.

### *S. mediterranea* shows age-related eye phenotypes that can be reversed by the process of regeneration

To study aging in planaria, we turned to *S. mediterranea* strains that proliferate via sexual reproduction, and defined zygotes as age zero. We found that most of these strains, including both wild isolates of unknown age and lines bred in captivity, develop visible morphological changes in the eyes between 6 months (6 MO) and 5 years of age. The most prominent phenotype is the development of ectopic eyes or ectopic eye pigment cells (**Fig. 1A-D**). This eye phenotype is also observed in isolates of a closely related species, *Schmidtea polychroa*, directly captured in the wild (**Fig. 1E**).

**Figure 1.**
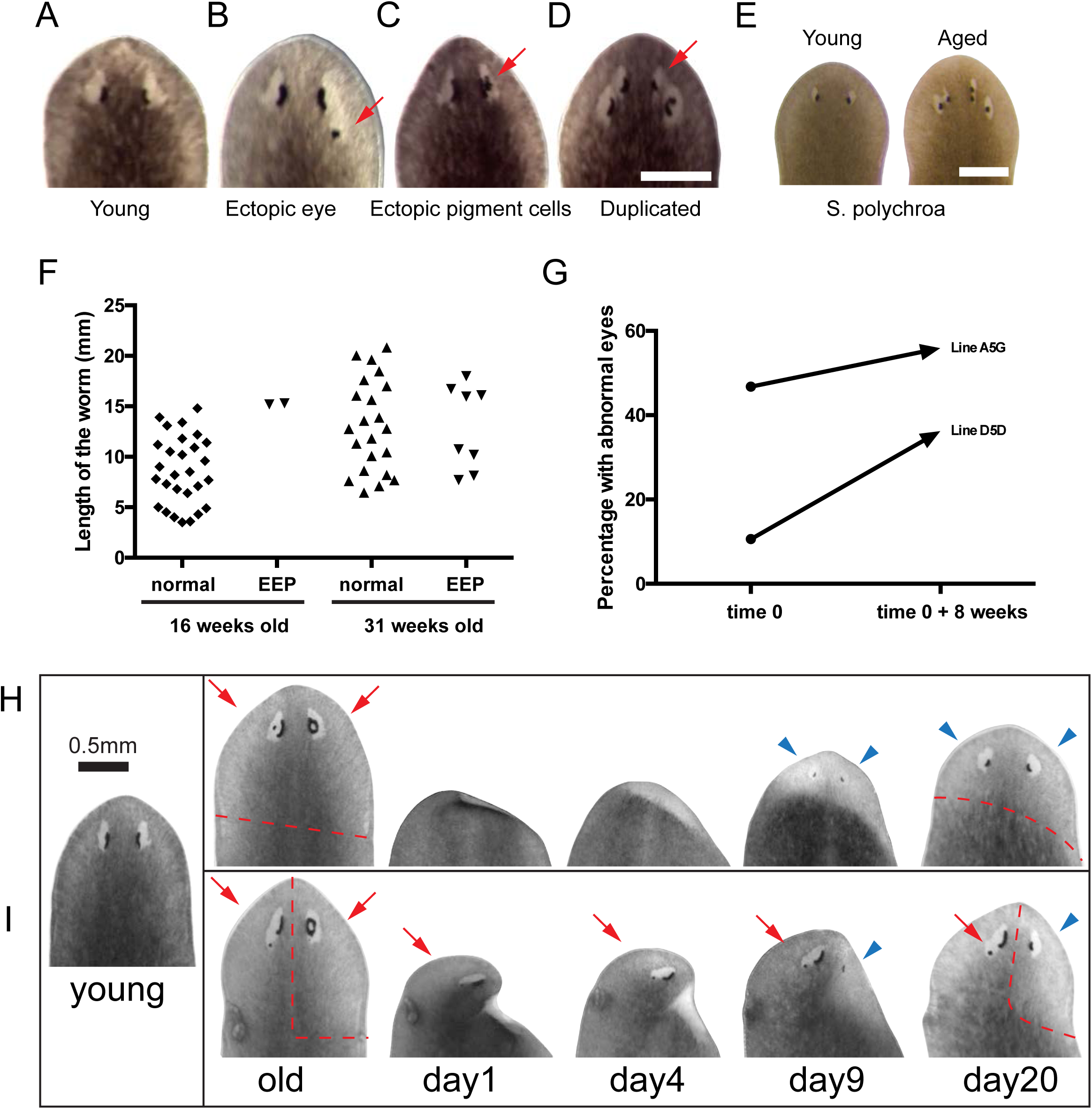
Aging and rejuvenation of planarian heads. (A) Young planarian heads, normal eyes. (B-D) Abnormal eyes in old planarians. Red arrow: eye pigment cells. (E) Abnormal eyes of wild-caught *S. polychroa.* Young *S. polychora* were born and bred in the laboratory as a control. (F) The same cohort of hatchlings were tracked and examined at the age of 16-week-old and 31-week-old. Each square and triangle represented one animal of the cohort. (G) Cohort sizes for line A5G and line D5D were both 50 animals each. (H-I) Original heads with two abnormal eyes, and regeneration after head removal (H) or half-head removal(I). Dashed line: amputation plane. Red arrows: old eyes. Blue arrowhead: new eyes. Scale bar = 0.5mm.

To confirm that the development of ectopic eyes or pigment cells (EEP) is age-associated, we performed two experiments. In the first experiment, we tracked a cohort of planarians from the time of their birth from egg capsules^40^. At the age of 16 weeks, when all animals were sexually mature, 2 out of 30 (6.7%) showed the EEP phenotype. At the age of 31 weeks, this fraction increased to 8 out of 30 (26.7%) (**Fig. 1F**). Notably, the development of EEP did not correlate with animal size (**Fig. 1F**). In a second experiment, we tracked clones of one individual (line D5D) that were produced by repeated amputation and regeneration. After this cohort of clones (n=50) reached sexual maturity, and at the first time point of observation, 10% of the animals showed the EEP phenotype. Eight weeks later, the fraction of animals with EEP increased to 35%. In a second cohort of clones (n=50) of a different individual (line A5G), we observed an increase of EEP from 46% to 55% after eight weeks (**Fig. 1G**). Thus, in several different experimental paradigms, the frequency of EEP phenotypes increases with age.

While generating clones for the above experiments, we noticed that amputating the heads of old planarians exhibiting EEP led to the generation of new heads lacking this phenotype. We wondered whether the amputation and regeneration process had somehow reset the age of the head. We therefore examined directly whether tissue loss and regeneration could reverse aging phenotypes. We removed the head from individuals with EEP phenotypes at the age of 32 MO and examined the newly regenerated heads. In these experiments we either removed the whole head (i.e., both eyes) to examine whether the remaining body would regenerate a head with two young eyes (**Fig. 1H**), or removed half of the head to examine whether the newly regenerated half of the head would have a young eye (**Fig. 1I**). In both experiments, amputation and regeneration produced young eyes. These young eyes developed EEP again in ~6 months post-regeneration.

To further characterize the rejuvenation effect and determine whether rejuvenation induced by repeated rounds of amputation/regeneration led to delayed aging, we compared the eye aging phenotypes of two cohorts of clones that were created from a single individual and were both chronologically 36 MO. The ancestral individual was obtained by mating a pair of adults (**Fig. 2A**). We repeatedly amputated and regenerated this single individual into a cohort of 181 clones, allowing three weeks for regeneration plus two weeks of feeding post-regeneration. We divided this cohort into a normal aging group (group A), which contained 91 clones and was maintained on a weekly feeding schedule for 30 more months, and a repeated-regeneration group (group B), which started with 90 clones, but underwent three additional rounds of amputation and regeneration over the same 30 months. During the expansion of group B, regenerated fragments (e.g., heads, trunks, tails) were mixed and fed once a week until they reached sexual maturity, at which point we initiated the next round of amputation and regeneration. At the end of the experiment, group B contained 2000 clones (**Fig. 2A**). We observed that 14 of 91 individuals in group A (15%) had developed EEP phenotypes, with sufficient heterogeneity in the phenotypes to distinguish one individual from another (**Fig. 2B**). If the two groups had the same aging rates, we would expect 15% of the clones in group B to develop EEP. Contrary to this expectation, a random sampling of 217 individuals from group B did not find any individuals exhibiting EEP (0%, the chi-square test, p<0.01) (**Fig. 2C**). All individuals in group B showed a uniform youthful eye morphology, suggesting that repeated amputation and regeneration induced rejuvenation of the eye and delayed the occurrence of the aging eye phenotypes.

**Figure 2.**
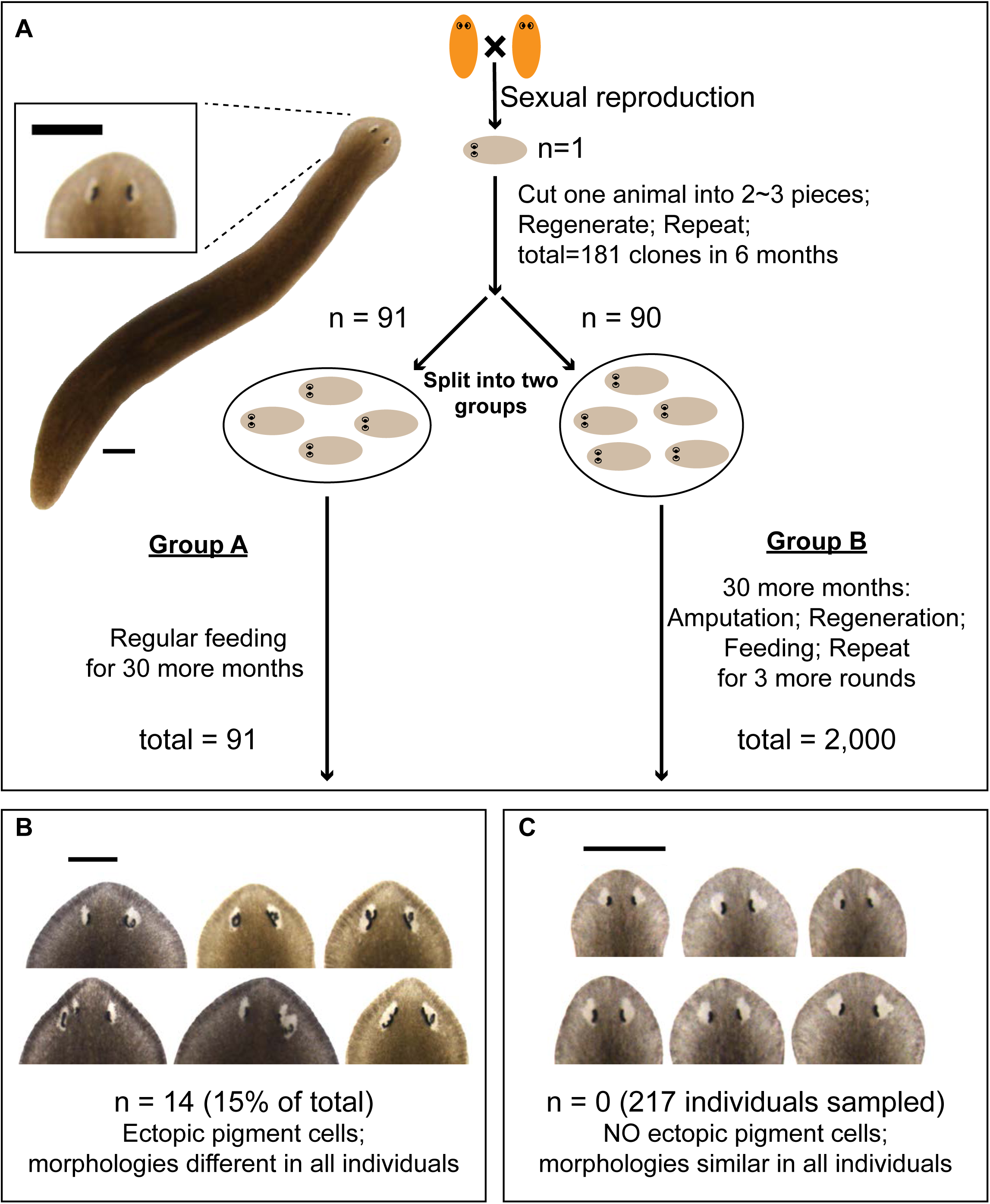
Repeated amputation and regeneration maintains youthful heads. (A) Theme of amputation, regeneration, feeding and experimental design. Scale bar = 1mm. (B) Representative images of 6 clones with heterogeneous EEP phenotypes from group A. (C) Representative images of 6 clones with homogenous young eyes from group B.

### Regeneration reverses age-related physiological phenotypes

To determine whether the rejuvenation effect was specific to the eye, or if it impacted other age-related phenotypes more globally, we examined additional physiological traits. A common age-related trait in many animals is reproductive senescence. We assessed planarian fertility by counting both the number of egg capsules produced and the number of hatched egg capsules every 100 days in cohorts of clones of an inbred line. We then measured fertility by tracking the percentage of egg capsules hatched over two years. We found that fertility declined continuously from ~40% at day 200 to ~10% at day 600 (**Fig. 3A, Supplementary Table 1**). Planarians older than two years continued to produce egg capsules, but their total number was reduced and <10% hatched, suggesting that reproductive senescence had occurred. We observed reproductive senescence in all sexual lines/populations maintained in captivity.

**Figure 3.**
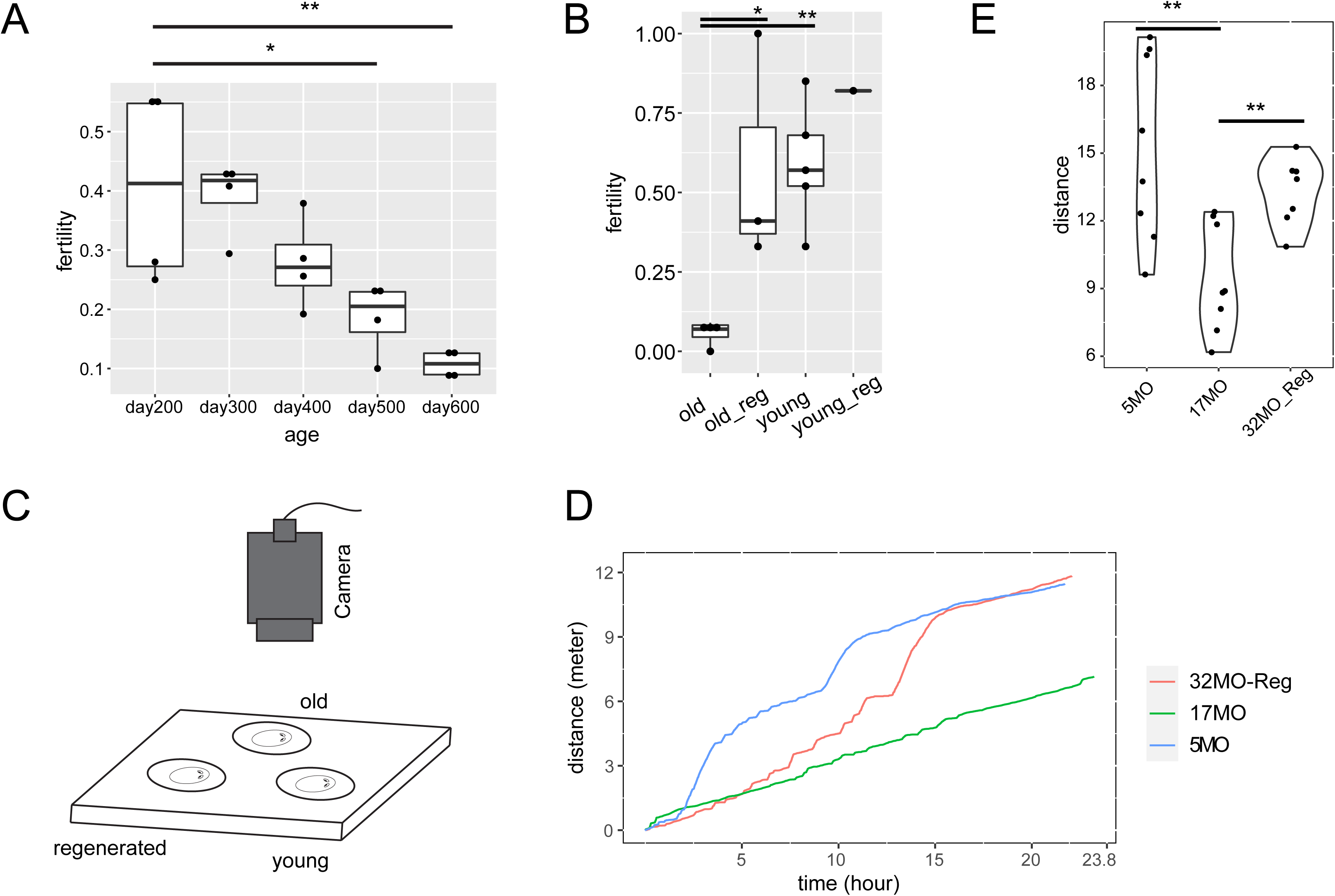
Aging and rejuvenation of fertility and motility. (A) Four biological replicates. Day 200 v.s. day 500, and day 200 v.s. day 600 were statistically significant, with p.values <0.05 and <0.01. One-Way ANOVA, with Tukey’s ‘Honest Significant Difference’ method. (B) Young cohorts: 4MO, n=1; 6MO, n=3, 7MO, n=1. Old cohorts: 12MO, n=1; 13MO, n=1; 14MO, n=1; 24MO, n=1. Old_reg cohorts: cutting old animals into multiple random fragments, n=1, 5 months post amputation of 24MO animals; cutting old animals into 9 fragments, n=1, 3 months post amputation of 18MO animals; removing the heads only, n=1, 3 months post amputation of 21MO animals. Old v.s. old_reg, and old v.s. young were statistically significant, with p.values < 0.05. One-Way ANOVA, with Tukey’s ‘Honest Significant Difference’ method. (C) Camera and chamber set up for video recording of planarian motility. (D) Distance traveled (in meters) over time (in hours) plot for one set of tested individuals. (E) Eight replicates. Each replicate measures distance traveled by one 5MO, one 17MO, and one 32MO-Reg individual. 32MO-Reg: A 32MO population was amputated at the age of 28MO. After regeneration, it was maintained for an additional 4 months before the experiment. Welch Two Sample t-test. p.values < 0.01 for 5MO v.s. 17MO and 17MO v.s. 32MO-Reg.

We then asked whether reproductive senescence can also be reversed by regeneration. We plotted fertility data for young, old and regenerated cohorts of animals from an inbred line. Fertility in the old cohorts (12-24 MO) was ~6% (**Fig. 3B**). Regeneration of the 18-24MO cohorts restored fertility to 58% (**Fig. 3B**), comparable to that of young (4-7MO) cohorts, which had an average fertility of 59% (**Fig. 3B, Supplementary Table 1**). Hence, we conclude that planarian regeneration also reverses age-related reproductive senescence.

Impaired motility is another characteristic of old age in mice and humans^41,42^. To test whether aged planarians exhibit impaired motility and, if so, whether regeneration can reverse it, we measured the total distance traveled by individual animals in 24 hours (**Fig. 3C-E, Supplementary Table 2**). We found that young (5MO) planarians were more active than older (17MO) planarians, traveling an average of 6 more meters in a day (**Fig. 3E**). The regenerated cohort (32MO-Reg) traveled an average of 4 more meters in a day than the 17MO individuals (**Fig. 3E**). We conclude that regeneration also reverses age-associated loss of motility.

### Single-cell transcriptomic analysis defines cell populations of the planarian head

Most animals exhibit age-related decline in stem cell function, which reduces regenerative abilities^2^. A decline in stem cell function could underlie the physiological effects we observed in aging planaria, yet the continued ability to regenerate may reflect maintenance of stem cell function. To understand the role of planarian stem cells during aging, we first characterized the types of stem cells found in the sexual lineage of *S. mediterranea*.

We used single cell mRNA sequencing (scRNA-seq) to profile head cells. To improve consistency of tissue sampling from different individuals, we used the posterior ends of the two auricles to mark the amputation sites and profiled each head with scRNA-seq. Head removal by this procedure leads to ~8.5% loss of body mass (**Fig. 4A**), minimizing the massive body remodeling that occurs in traditional planarian amputation experiments^24,43,44^. New heads are regenerated within 20 days, with no significant changes in whole body shape and size, allowing us to consistently analyze tissue types from different chronological ages and from different animals that underwent perturbation by regeneration.

**Figure 4.**
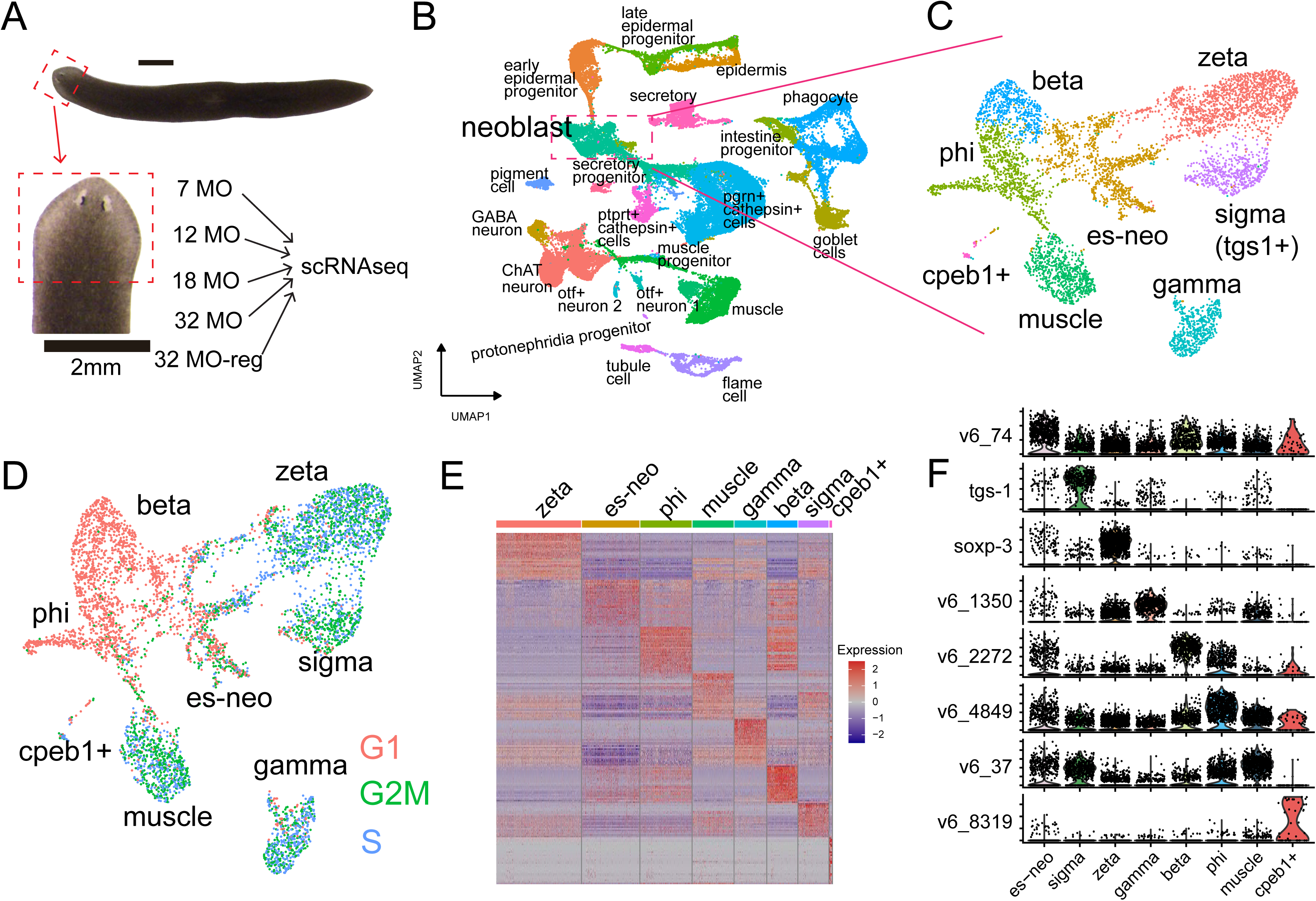
Single cell profiling of the heads and the sexual adult stem cell system. (A) Head amputation and chronological ages of the samples. Young: 7 MO (n=2), Aged 12 MO, 18 MO, 32 MO (n=1 each), Reg: 32 MO-regenerated (n=3). (B) Major cell types identified from the heads. (C) Subclusters of sexual adult stem cells (neoblasts). (D) Cell cycle distribution for each type of adult stem cells. (E) Gene expression heatmap for each type of adult stem cells. Gene expression in es-neo is most stochastic. (F) A significant number of es-neo cells express high levels of markers representing every other stem cell type. Dots represent individual cells. y-axis represents expression levels. Gene names were from previous studies (tgs-1^26^, soxp-3^47^) or dd_Smed_v6^80^ (v6_numbers).

We obtained a total of 53,136 high quality single cell transcriptomes from heads at different ages. The heads were obtained from individuals that were 7 MO (2 heads), 12 MO (1 head), 18 MO (1 head), and 32 MO (1 head). In addition, regenerated heads were obtained from three individuals which were amputated at the age of 32 MO and allowed to regenerate for 2 months (**Fig. 4A, Extended Data Fig. 1a,b, Supplementary Table 3**). Unsupervised clustering of the data identified cells from all eight major planarian tissue types^45,46^, including neoblasts (planarian stem cells), epidermis, secretory cells, intestine, protonephridia, cathepsin+ cells, neurons, and muscles (**Fig. 4B, Extended Data Fig. 1c,d**). With unsupervised sub-clustering of these eight tissue types, we identified a total of 97 sub-clusters (**Fig. 4C, Extended Data Fig. 2**), and annotated those from neoblast, neurons, and muscles (**Supplementary Table 4**).

We then focused on the stem cell populations. In general, we found that the adult stem cell system of sexual planarians is comparable to that of asexual planarians, which contains multiple stem cell subtypes (**Fig. 4C**). Sexual adults harbored several stem cell subtypes that appeared homologous, based on their transcriptomic signature, to stem cells in asexual animals, and hence likely have similar differentiation potentials. These include sigma (soxP-1^+^, soxP-2^+^), zeta (soxP-3^+^, zfp-1^+^, egr-1^+^, p53^+^) and gamma (hnf4^+^, gata456-1^+^, nkx2.2^+^) neoblast populations^47,48^ (**Fig. 4C, Supplementary Figure 1**). Indeed, an analysis of cell cycle states associated with these cells was consistent with this idea. Previous work has shown that stem cell differentiation is associated with cell cycle progression^49^. Consistent with these findings, we found that progressions from zeta neoblasts to early epidermal progenitors (p53^+^, prog-1^+^)^47,48,50^, and from gamma neoblasts to intestinal cells (apob-1^+^ or apob-2^+^)^51,52^, were both associated with a transition from the S/G2/M phases to the G1 phase (**Supplementary Figure 2-3**). We confirmed cell cycle states by examining additional markers of proliferation (mcm4, mcm5, mcm7) and post-mitotic cells (xbp-1,p4hb)^49^ (**Supplementary Figure 2-3**). Furthermore, we found that the pluripotent tgs-1^+^ cells^26^ are sigma neoblasts. In the sexual planarians, ~58.8% (+/− 2.6%, SEM) of tgs-1^+^ cells are soxP-1^+^ and soxP-2^+^ (sigma neoblasts), while 75.0% (+/−2.7%, SEM) of sigma neoblasts are tgs-1^+^. Tetraspanin-1, the antibody of which was used to purify tgs-1^+^ adult pluripotent stem cells in asexual planarians^26^, correspond to 26.2% (+/− 2.0%, SEM) of sigma neoblasts (38% of tspan-1^+^ cells, +/−3.4%, SEM) and to 19.2% (+/− 2.1%, SEM) of muscle progenitors (37.2% of tspan-1^+^ cells, +/−3.7%, SEM) (**Supplementary Figure 4, Supplementary Table 5**).

Sexual planarians also harbor four previously uncharacterized stem cell types, which we name phi, beta, CPEB1+ and es-neo neoblasts. Among the phi and beta cells, which account for 1.7% and 1% of the head mass, respectively, 89.7% (+/−1.6%, SEM) and 99.3% (+/−0.3%) of the cells, respectively, are in the G1 phase (**Fig. 4D, Supplementary Table 5**), similar to the pattern in early epidermal progenitors (prog1+ cells)^48^. Both phi and beta neoblasts express high levels of foxF-1 (**Supplementary Figure 5**), a gene required for the differentiation of dorsal-ventral muscles (DVM), intestinal muscles (IM), and cathepsin+ parenchymal cells in asexual planarians (**Supplementary Figure 6**)^53^. These cells also express high levels of *F-spondin* ^54^, *LDLRR-3*^54^, and ETS-1^55^ (**Supplementary Figure 9**). Phi neoblasts show somewhat higher expression of *foxF-1*, *nf1c* (dd_v6_12312) and *ptprj* (dd_v6_4490), and lower expression of *F-spondin* and *LDLRR-3* (**Supplementary Figure 10**), which suggest that they may be precursors to beta neoblasts.

Several lines of evidence suggest that phi and beta cells are progenitors of cathepsin+ parenchymal cells, rather than muscle cells. First, phi and beta neoblasts do not express muscle-differentiation-related transcription factors, such as myoD and nkx1-1^56^, which are normally expressed in muscle progenitors (**Supplementary Figure 7**). On the other hand, they do show enriched expression of cathepsin+ cell markers^57^ (**Supplementary Figure 8**). Furthermore, *F-spondin* ^54^, *LDLRR-3*^54^, and ETS-1^55^ have all been associated with parenchymal cells: *F-spondin* and/or *LDLRR-3* are secreted by groups of differentiated and irradiation resistant brain parenchymal cells in asexual planarians^54^, and loss of *ETS-1* function has a specific effect on gene expression in cathepsin+ cells^55^. Finally, differentiation trajectory analyses by CytoTRACE^58^, based on mRNA quantity and diversity, CellRank^59^, which uses transcript similarity and mRNA splicing dynamics computed by RNA velocity^60^, and SlingShot^61^, which computes a minimum spanning tree and simultaneous principal curves from clusters defined by transcript similarity, further support this conclusion (**Extended Data Fig. 3e-g**).

CPEB1+ neoblasts only accounted for 0.08% (+/−0.03%, SEM) of the head mass (**Supplementary Table 5**). The top genes enriched in this group (**Supplementary Figure 11**), including tyrosinase, C-type lectin-like protein, capsule tanning factor 1 and surfactant B-associated protein, are all expressed in yolk glands, and are required for tanning of the egg capsule shells^62^, suggesting that this group of neoblasts represents progenitors of yolk glands. The high and specific expression of CPEB1 (**Supplementary Figure 11**) supports this conclusion, given that knockdown of CPEB1 leads to loss of yolk glands and ovaries^62^. Due to the small number of CPEB1+ neoblasts, their aging and rejuvenation process were not examined in depth.

Es-neo stem cells accounted for 1.9% of the head mass (+/−0.7%, SEM, **Supplementary Table 5**). This group of neoblasts had three unique features. First, es-neo is the only group of stem cells with comparable proportions of S (17.9%, +/−2.2%, SEM), G2/M (23.1%, +/−1.5%, SEM), and G1 (59%, +/−3.4%, SEM) cells (**Fig. 4D, Supplementary Table 5**). The rest of the sub-types either lack a G1 phase (i.e., actively dividing, sigma, zeta, gamma, and muscle) or are dominated by G1 (i.e., post-mitotic, phi and beta) (**Fig. 4D**). Second, es-neo cells show high expression of a range of canonical markers, including general markers for stem cells (e.g., smedwi-1, 2, 3), for sub-types of stem cells (e.g., sigma, zeta, gamma, **Fig. 4F**), and for specific differentiated lineages (e.g., epidermis, muscle, intestine, protonephridia, nervous system) (**Supplementary Figure 12**). Finally, the expression of these marker genes is highly stochastic (**Fig. 4E-F**); the cells do not show consistent or deterministic marker expression, and no markers are expressed in 100% of es-neo cells. As a result, no unique marker genes were identifiable for es-neo. The es-neo cluster is defined by the stochastic state of gene expression, which is not affected by cell cycle transition. These gene expression patterns are reminiscent of the stochastic expression of pluripotency factors in embryonic stem cells (e.g., Nanog, Oct4) (reviewed by MacArthur and Lemischka^63^), a likely signature of pluripotency. Trajectory analysis with CellRank^59^ (mRNA splicing), and Slingshot^61^ (transcript similarity) both support the idea that es-neo may be a totipotent stem cell population that gives rise to zeta, sigma, phi, muscle, gamma and CPEB1+ cells (**Extended Data Fig. 4**).

### Age-dependent gain and loss of specific cellular populations can be reversed by regeneration

We next asked how aging affects stem cells and other cell types of the head. In mammals, most tissues, with the exception of the hematopoietic system^2^, exhibit an age-dependent decline in the total number of stem cells. In our experiments in sexual planaria, however, we did not observe a significant loss of stem cells with age (**Fig. 5A, Extended Data Fig. 5**). Instead, there was a significant increase in the number of beta stem cells in older animals, and we also observed an upward trend in the number of zeta stem cells as well between 7 MO and 32 MO. The lack of statistical significance for the latter observation is likely due to the smaller sample size, or the fact that the increase becomes extensive only in much older individuals (>18 MO) (**Extended Data Fig. 5b**).

**Figure 5.**
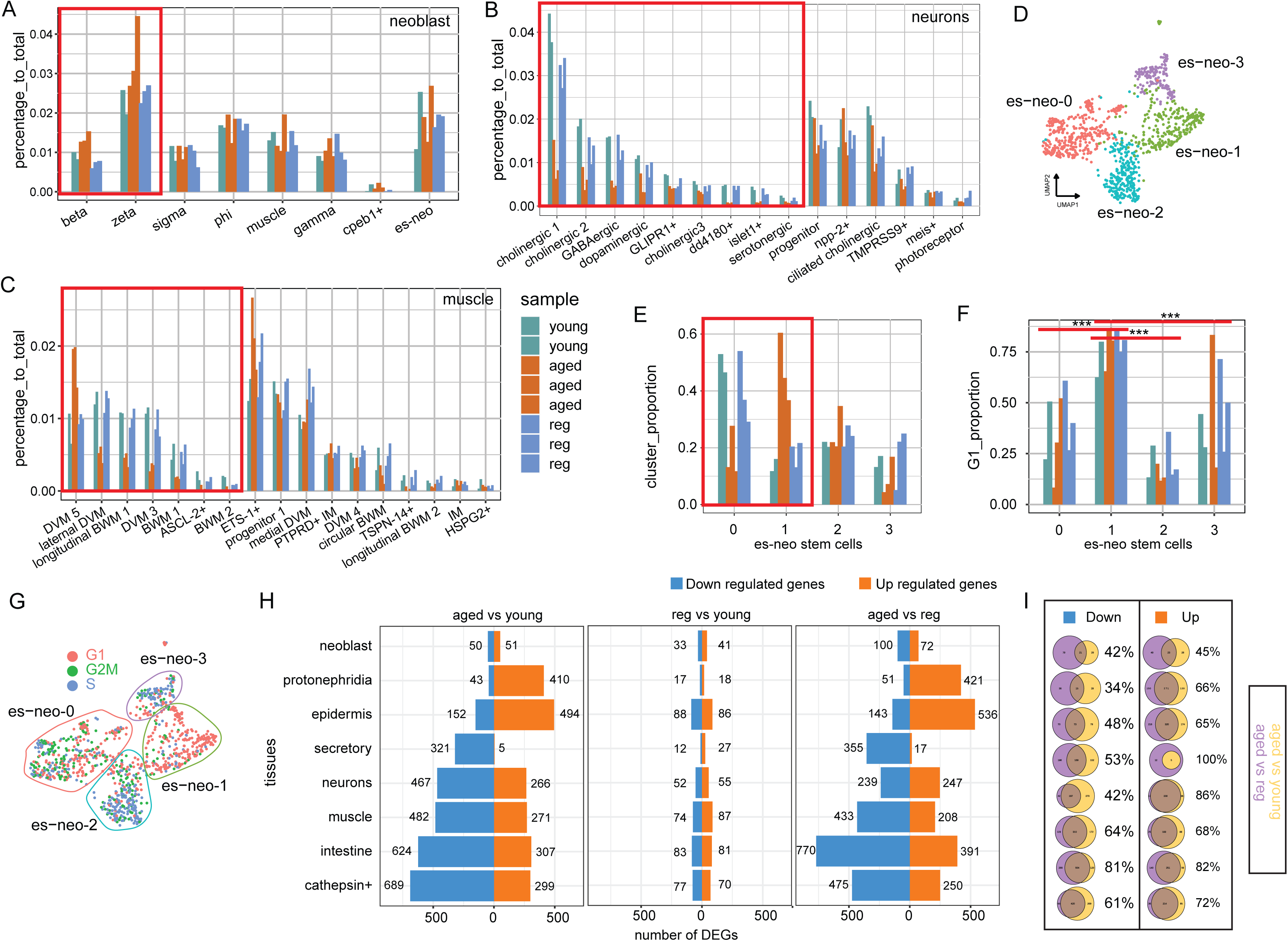
Cellular and molecular rejuvenation of aged planarians. (A-C) Proportion of each cell type to the cells in the whole head. (A) stem cells. (B) neurons. (C) muscles. Young: 7MO. Aged:12MO, 18MO, 32MO. Reg:32MO-regenerated. Red rectangle: cell types that are significantly reduced/increased in older planarians, and significantly reversed to young after regeneration. p.values < 0.05. One-Way ANOVA, with Tukey’s ‘Honest Significant Difference’ method. (D) Four distinct subclusters of es-neo stem cells. (E) Within es-neo population, es-neo-0 is significantly reduced in older planarians, while es-neo-1 is significantly increased. (F-G) Cell cycle distribution in four es-neo subclusters (G). G1 cells of es-neo-1 is significantly more than G1 cells of three other subclusters, p.values < 0.01, One-Way ANOVA, with Tukey’s ‘Honest Significant Difference’ method (F). (H) Number of differentially expressed genes between aged and young (left), regeneration and young (middle), and aged and regeneration (right), in eight tissue types. Blue=down regulated genes. Orange=up regulated genes. (I) The exact same genes shared by aged vs reg, and aged vs young, and relative percentage of such number of genes to aged vs young. Down regulated genes (left). Up regulated genes (right). Each row corresponds to the tissue type in H.

By contrast, and consistent with aging in mice and humans, planarian aging is also accompanied by a drastic loss of muscle cells and neurons (**Fig. 5B-C**), which may contribute to the loss of fertility and motility (**Fig. 3**). Compared to young (7 MO) planarians, old (12MO, 18MO and 32MO) planarians, lost 67.9% of the neurons in 9 out of 15 neuronal cell types, which dropped from 11.1% (+/−0.3%, SEM) to 3.6% (+/−2.0%, SEM) of the head mass (**Fig. 5B, Supplementary Table 5**). The other 6 neuronal types remained unchanged. In the muscle, 6 out of 17 cell types lost 65.3% of the cells, dropping from 4.4% (+/−0.16%, SEM) to 1.5% (+/−0.9%, SEM) of the head mass, although one dorsal-ventral muscle population (mus-2) increased by 2.08-fold, from 0.8% (+/− 0.2%, SEM) to 1.8% (+/−1%, SEM) of the head mass. The other muscle cell types did not show a significant change. Overall, there was a net 2% loss of muscle mass in the head (**Fig. 5C, Supplementary Table 5**).

Other major tissue types (i.e., epidermis, cathepsin+ parenchymal cells, protonephridia, secretory cells), showed a mixed pattern, in which planarian aging was accompanied by both gain and loss of specific cell types^4^. In the protonephridia system^64–66^, aging led to loss of 66.7% of tubule cells (pro-1) and 58.8% of flame cells (pro-6), but a two-fold expansion of the collecting duct cells (pro-3) (**Extended Data Fig. 3a-c, Supplementary Table 5**). In the cathepsin+ cell system, trajectory analysis with CytoTRACE, CellRank and SlingShot all support the idea that ldlrr-1+ cells (cat-1/cat-3 and cat-2) are differentiated progeny of progenitor cells (cat-0) (**Extended Data Fig. 3d-g**). Aging led to a ~2.5-fold expansion of cat-1 and cat-3 cells, but a 44.8% loss of cat-2 cells (**Extended Data Fig. 3d, Supplementary Table 5**), suggesting that the differentiation potential of cat-0 is likely shifted towards cat-1/cat-3 with age, a phenomenon similar to aging in human hematopoietic stem cells^67^. In the epidermal system, aging led to an expansion of early epidermal progenitors (epi-0, epi-2), but a loss of late epidermal progenitors (epi-5) and mature epidermis (epi-8) (**Extended Data Fig. 3h-k, Supplementary Figure 13**).

For es-neo stem cells, there was no change in the total numbers, but we observed a shift from the S/M cell cycle phases to G in 32 MO planarians (**Fig. 5D-G**). Unsupervised sub-clustering of the heterogeneous es-neo populations showed four distinct sub-populations, es-neo-0 to es-neo-3 (**Fig. 5D**). As planarians aged, the proportion of the es-neo-0 population decreased, accompanied by a significant increase in the es-neo-1 population (**Fig. 5E**). Most es-neo-1 cells are not in the S or M phases (>77%, **Supplementary Table 6, Fig. 5G**), and the proportion of these cells in aged animals is significantly higher compared to the other three subclusters (**Fig. 5F**), which may explain the shift in cell cycle states.

We then asked whether these cell-type specific changes could be reversed by regeneration. Indeed, single cell transcriptomic analysis showed that regeneration reversed the age-dependent changes in beta/zeta stem cell, muscle and neuron cell numbers, as well as shifts in es-neo stem cell cell cycle states, restoring the youthful composition and character of all the cell types (**Fig 5A-F, Extended Data Fig. 3 and Supplementary Table 5**).

### Regeneration reverses aging-dependent molecular changes

Finally, we sought to understand the molecular changes that underlie rejuvenation. To do so, we examined changes in gene expression between young, old and regenerated planarian heads. All tissues showed changes under these different conditions, but the degree and pattern of gene expression changes varied. Comparing old and young planarians, for example, the neoblast compartment, had fewer differentially expressed genes (DEGs) relative to differentiated tissues (**Fig. 5H-I**). Furthermore, neoblasts showed a balance between genes that were down- and up-regulated (50 downregulated genes versus 51 upregulated, in aged vs young planarians). In protonephridia and epidermis, on the other hand, many more genes were up-regulated in aged planarians, whereas in secretory cells, the nervous system, muscles, intestines and cathepsin+ cells, many more genes were down-regulated in aged planarians (**Fig. 5H**). The tissue specific pattern of DEGs suggests that aging has tissue-specific impacts. Notably, the pattern of gene expression changes between aged and young planarians was fully recapitulated when we identified DEGs in aged v.s. regenerated planarians (**Fig. 5H, right**). These results suggest that regeneration led to a global rejuvenation of gene expression in all tissues.

In further support of this conclusion, comparing young and regenerated planarians revealed only a small number of DEGs (**Fig. 5H, middle**). In fact, when looking at individual genes showing either an increase or a decrease in expression, we find that regeneration led to upregulation of 34-81% of the genes that were downregulated in aged individuals relative to young ones in different tissues (**Fig. 5I, left**); conversely, regeneration led to downregulation of 45-100% of genes showing upregulation in aged individuals in different tissues (**Fig. 5I, right**). Thus, regeneration reverses most age-dependent changes in gene expression.

Beyond the general patterns of gene expression shifts, the specific molecular changes revealed by our gene expression analysis indicate parallels in the effects of aging between sexual planarians and traditional animal models^1^, and here, too, there are tissue specific effects^68^ (**Extended Data Fig. 6 and 7**). For example, we observed downregulation of insulin and target of rapamycin complex-1/2 signaling in the nervous system (neu-2, npp-2+ neurons), in muscle (mus-0, ETS-1+ cells), and in the cathepsin+ parenchymal cells (cat-4, psap+ cells). By contrast, we found alterations in the expression of genes regulating mitochondrial functions in all cell types. These results highlight conserved aspects of the aging process, yet underline cell-specific patterns.

The fact that some cell types show many age-dependent gene expression changes, while other cell types show only a few, suggests that the tissues may exhibit differential vulnerability to age— i.e., that cell types showing a larger number of changes may be more vulnerable to the effects of aging (**Extended Data Fig. 7, Supplementary Table 7**). Our results suggest that the number of changes—and hence vulnerability to aging—may correlate with the extent of differentiation. For example, in the nervous system, npp2+ neurons (neu-2) were most sensitive to age-induced transcriptional alteration, showing changes in the expression of 473 genes. Ciliated cholinergic neurons (neu-3) and neuronal progenitors (neu-1), which were similarly abundant to neu-2 neurons in the heads (~2%, **Fig. 5B**), showed changes in only 91 and 25 genes (**Extended Data Fig. 7**). We propose that cells that are less differentiated (medium CytoTRACE values) are more sensitive to age-induced transcriptional alterations than highly pluripotent (high CytoTRACE values) or highly differentiated cells (low CytoTRACE values) (**Supplementary Table 7**).

Our studies reveal a naturally evolved solution to age reversal in planarians. We show that rejuvenation reverses the effects of aging on multiple tissues, hundreds of genes, and several physiological traits. Considering the life history of these animals is near “immortal”, the extent of rejuvenation is likely more comprehensive and happens on the whole-body scale. Understanding the regulatory mechanism of global rejuvenation has the potential to point the way toward new classes of anti-aging interventions.

## Methods

### Planarian strains, lab culture and fertility

All planarians were maintained in 1x Montjuïch salts, in plastic containers, at 18°C, with once a week feeding of organic cow liver as previously described^69^. All animals used in the current study were sexually mature and starved for a week before experimentation. Fertility data was collected by documenting the number of egg capsules produced every week by the examined cohorts. The number of hatchlings and the number of hatched egg capsules were documented for each batch of egg capsules 3 weeks after collection. Fertility value was calculated as the percentage of egg capsules that hatched divided by the total number of egg capsules collected over a defined period.

Inbred strains used in current study include A5G, D5D, D2E, S2F8b, S2F12 and LAF. Each assay reported has been repeated in multiple inbred strains, instead of using one single strain or a mixed population with drifting genetic backgrounds.

### Single cell RNA Sequencing Sample Preparation

Heads of the animals were surgically removed under a dissection microscope, along the posterior end of the auricle. Tissues were macerated with a surgical knife in CMF buffer supplemented with 0.5% BSA (CMFB) as previously described^26^. Macerated tissues were kept on ice-cold CMFB in a 6-cm petri dish with gentle horizontal shaking for 20min. Tissues were pipetted up and down every 5 minutes with a transfer pipette for 5 to7 times to ensure sufficient disassociation and optimal viability. Cells were filtered through a 40 μm strainer before collection by spinning at 400rcf for 5min at 4°C. Cells were counted manually. Cell viability was assessed by labeling with trypan blue (0.4%). A general viability reading of more than 70% is required before library preparation with 10x single cell 3’GEM kit. To compare the young, aged, and rejuvenated heads, single cells were prepared on the same day for one head from each condition. Libraries were prepared and sequenced in the same batch. In general, each cell was sequenced to 35k to 80k reads on NovaSeq S4 platform.

### Single-cell gene expression quantification and data quality control

Using CellRanger software V6.0.1 (10x Genomics), the sequencing data were pre-processed and mapped to the reference genome^29,30^, and the count pipeline was run to generate the gene expression matrix. Low-quality cells were filtered to remove signals from empty droplets or degraded RNA by discarding cells with less than 500 detected genes, and cells with high expression levels of mitochondrial genes (**Supplementary Table 3**). Doublets were detected by identifying outliers with extremely high detected UMI numbers (**Supplementary Table 3**) and by employing DoubletFinder^70^ algorithm and were removed from each sample (**Supplementary Table 3**).

### Data integration, clustering, and visualization

Totally 53,136 cells from 8 samples were kept for subsequent analysis. The UMI counts of each cell were normalized by Seurat^71^, and the normalized data were log-transformed for further analyses and visualization. The top 2,000 genes with the highest standard deviations were obtained as highly variable genes (HVGs). For the integration of different condition samples, Seurat anchor-based method^72^ was performed to identify shared sources of variation across datasets, allowing for robust identification of cell types and differential gene expression analysis. Dimension reduction using the PCA method was performed, and the top 30 PCs were used for downstream clustering analysis. The graph-based clustering approach was used for cell clustering with the resolution parameter set to 0.4, resulting in 32 distinct cell clusters. Uniform Manifold Approximation and Projection (UMAP) was used for visualizing cell distance in reduced two-dimensional space.

### Cell cluster annotation and subclustering

To annotate the clustering results of the whole dataset to specific tissue types, two steps were followed. (1) The cluster markers of each cluster were calculated, and clusters expressed classic markers of known tissue type were assigned to the same tissue type. (2) The expression patterns of clusters were compared to the comprehensive planarian cell atlas^45,46^, clusters that shared high similarity with certain cell types were considered to be the same tissue type. Cells of each tissue type were subset from the whole dataset and integrated following the similar processes mentioned above. After significant PCs were identified (1-20 PCs were used for subclustering of cells in each tissue type), graph-based clustering was performed with variable resolution parameters according to the tissue heterogeneity. For some tissues, for example, neoblast, the sub-clusters were annotated as specific cell types according to the expression of cell-type specific markers. UMAP was used for sub-cluster visualization.

### Cell cycle analysis and CytoTRACE analysis

For each tissue type, the cell-cycle state (G1, G2/M, S) of cells in the data was computationally predicted by the expression of cell-cycle-associated markers^73^ (S and G2M, **Supplementary Table 9**). The CellCycleScoring function in Seurat was performed to assign the S and G2M cell cycle score to each cell, along with the phase assignment in either G1, G2M, or S phase based on the cell cycle score. CytoTRACE^58^ was applied to order cells by their differentiation status using default parameters for each single tissue type. Cells with higher CytoTRACE scores are less differentiated and with higher developmental potential, and the verse visa.

### Differential gene expression analysis and gene ontology enrichment analysis

Differential gene expression analysis was conducted to characterize gene **expression** changes during aging and after regeneration. For each tissue type and sub-clusters, differential expression genes (DEGs) between young and aged, aged and regenerated, and young and regenerated pairs were identified using FindMarkers function^71^ which employs Wilcoxon rank sum test, with p-values adjusted by the Benjamini-Hochberg method for multiple testing correction. Significant DEGs were identified with an average |log2FC| > 0.25, and an adjusted p-value < 0.05. Gene ontology (GO) enrichment analysis of significant DEGs was performed using the EnrichPipeline^74^, and significantly enriched GO terms were identified with a false discovery rate cut-off of 0.05.

### Lineage and trajectory inference

The specialized neoblast subtype was combined with a specific tissue type to characterize the differentiation relationships. Slingshot^61^ algorithm (Version 2.6.0) was used to compute the single-cell pseudotime trajectory. Specifically, after dimension reduction and clustering steps in the Seurat analysis pipeline, the slingshot function was called to determine the lineage structure by constructing a minimum spanning tree (MST) between the clusters. Cell fate prediction was performed utilizing the following computation tools: velocyto^60^ (version 0.17.17) was used to quantify spliced and unspliced mRNA counts, scVelo^75^ (version 0.2.5) was used to compute RNA velocity vectors based on splicing ratios using the steady-state model on a per cell basis, and CellRank^59^ (version 1.5.1) was used to compute the initial and terminal states, which combines the robustness of trajectory inference with directional information from RNA velocity.

### Motility measurements

An automated video tracking setup was used to record the movements of animals^76,77^. Video recordings were taken for 24 hours and analyzed with the video analysis software DDrop^76,77^. Based on the tracking data, the cumulative distances, and total distances of each animal in 24 hours were calculated.

### *In situ* Hybridization Chain Reaction (HCR)

HCR probes for target genes were synthesized by Molecular Instruments. HCR RNA fluorescence *in situ* hybridization was carried out as previously described^30,78^. Briefly, the animals were treated by 10% NAC for 10-15min and fixed in 4% PFA for 20min. After washes with PBS + 0.5% Triton X, the animals were treated by Proteinase K for 20min at room temperature. Prehybridization, hybridization and amplification of HCR probes were carried out following standards of HCR protocol. Probes were used at 16nM. Hairpins were used at 60nM. DAPI was used to stain cell nuclei at the second last wash step. Samples were mounted with Prolong Diamond (Thermo Fisher) and procured for 24 hours before imaging.

### Microscope and Image process

Images were taken with Nikon A1 confocal using 10x, 20x or 40x lens. Images were read into Fiji (version 2.9.0/1.54d)^79^ or Imaris (version 9.9.1) and converted into RGB format. Quantification of cells were carried out manually in a double-blinded fashion.

## Acknowledgements

We thank Emma Belfer, Emily Warda, Brittany Pakfar, Clarice Chui, Jacob Fitz, Nabeel Karapatsakis for their assistance in planarian colony maintenance and breeding. We thank Drs. Safa Beydoun, and Joe Endicott for their help in assay establishment and exploration. We thank Florence Jinhao Guo and Athena Jinyue Guo for their assistance in the quantification of fertility. We thank Dr. Alejandro Sánchez Alvarado for his maintenance of original planarian inbred strains and generous support for our work. We thank Dr. Daniel Michele for his careful reading of our draft and insightful comments for revision. We thank Dr. Richard Miller for insightful and inspiring discussions on our experimental designs, data presentation and feedbacks on drafts of the manuscript.

## Contributions

LG: funding, conceptualization, experimental design, data acquisition and interpretation, draft, revision. XD: experimental design, data acquisition and interpretation, revision. XL, DP, JS: data acquisition. LK: funding, experimental design, revision. SP: data acquisition, revision.

## Figure legends

**Extended data Fig.1 Single-cell RNA sequencing data quality and cluster characterization a**, UMAP plot depicting the distribution of cells from different replicates. **b**, Violin plots of the number of UMIs (top) and genes (bottom) of each sample. **c**, UMAP plot of the entire dataset (n = 53,136) with cells labeled and colored by clusters (n = 32). **d**, Heatmap of the scaled expression of tissue markers, which assigned the clusters to specific tissues.

**Extended data Fig.2 Subclustering of eight planarian tissues**

UMAP plots of different tissues colored by their sub-clusters. **a**, neoblast (8 clusters, 5,751 cells). **b**, muscle (17 clusters, 5,571 cells). **c**, neurons (15 clusters, 7,092 cells). **d**, cathepsin+ parenchymal cell (11 clusters, 13,535 cells). **e**, epidermis (11 clusters, 7,183 cells). **f**, intestine (10 clusters, 8,226 cells). **g**, protonephridia (10 clusters, 2,497 cells). **h**, secretory (15 clusters, 3,281 cells).

**Extended data Fig.3 Aging and Rejuvenation in differentiated tissues**

**a-c**, Protonephridia. (**a**) Cell proportion of each sub-cluster in samples of different ages. (**b**) Lineage structure estimated by Slingshot. (**c**) UMAP plot showing piwi1^+^/piwi1^−^ protonephridia cells. **d-g**, Cathepsin^+^ parenchymal cell. (**d**) Cell proportion of sub-clusters. (**e**) Trajectory inferred by CellRank/RNA-velocity analysis with subpopulations of phi and beta included. (**f**) lineage structure predicted by Slingshot with phi and beta included. (**g**) UMAP plot showing CytoTRACE score of cells in different sub-clusters, with phi and beta included. The color indicates the degree of differentiation from low (red) to high (blue). **h-k**, Epidermis. (**h**) Trajectory inferred by CellRank/RNA-velocity analysis with subpopulations of zeta included. (**i**) Cell proportion of sub-clusters. Trajectory inferred by Slingshot (**j**) and CytoTRACE (**k**) of epidermis cells, with zeta included. Cell proportion of intestine (**l**) and secretory (**m**) sub-clusters.

**Extended data Fig.4 Cell differentiation trajectory of neoblast subclusters**

**a**, Differentiation trajectory by CellRank/RNA-velocity. UMAP of neoblasts with scVelo-projected velocities shown as streamlines. Clusters are colored and labeled according to original cluster annotations. **b**, Differentiation trajectory by Slingshot. A minimum spanning tree (MST) between neoblast cell types (nodes) was constructed by performing Slingshot algorithm to infer lineage trajectories.

**Extended data Fig.5 Estimation statistics of stem cell loss in old planarians**

Error bars on estimation plots show 95% Confidence Intervals for the differences between the means of the two tested age conditions (x-axis). There is 95% confidence that differences between regenerated and aged, between young and aged are in a tight range below 0 for beta stem cells (a). In other words, the significant increase of beta stem cells in older age (Fig 5A) is unlikely going to be wrong even if sample sizes are increased. The likelihood is less than 5%. On the other hand, the mean differences between age groups can be 0 in zeta (b), or most likely 0 in es-neo (c) and sigma (d) stem cells. For es-neo and sigma, increasing sample sizes is unlikely to change the conclusion that these stem cell types did not change between different age groups.

**Extended data Fig.6 Gene ontology of age-related transcriptional changes**

Bar graph (left) depicting number of differentially expressed genes between aged and young heads, and box plot (right) displaying the enriched Gene Ontology (GO) terms in different cell types. Orange: up regulated genes. Blue: down regulated genes.

**Extended data Fig.7 Molecular change of sub-clusters in all tissues with age**

Bar plot showing the number of differentially expressed genes in each sub-cluster between young (7 MO x2) and aged (12 MO, 18 MO, and 32 MO) heads. Orange bars represent up-regulated genes of aged heads to young heads, and blue bars represent down-regulated genes.

**Extended data Fig.8 Self-injury and fragmentation events**

**a**, A normal planarian. **b-d**, half-head injured worms. **e-h**, whole-head injured worms. **i-l**, Worms recovering from head-injury. The white regions indicate the blastema. **m**, pharynx-injured worm. **n-q**, head-fragments from pharynx-injury. **r-u**, tail-fragments from pharynx-injury. **v-w**, Worms with nearly complete regeneration. **x-y**, worms injured at copulatory apparatus. **z**, anterior and posterior-fragment from a copulatory apparatus-injury event.

**Supplementary Table 1 Aging and rejuvenation of sexual reproduction**

Table 1.1, Fertility rates at different ages in strain S2F12. Four independent repeats. Fertility = hatched eggs/total eggs. Population size for each repeat > 20. For each time point, fertility was calculated with accumulated egg data over 100 days.

Table 1.2, Fertility rates in young, aged and regenerated populations.

**Supplementary Table 2 Aging and rejuvenation of motility**

Total distances traveled by nine sets of different animals recorded in 24 hours. Young = 5MO. Aged = 17MO. Regenerated = 4 months after amputation of 28MO animals. Distance unit = meter.

**Supplementary Table 3 scRNA-Seq data summary**

Sample information, sequencing information, and quality control thresholds of each sample. you1/you2 = 7MO heads, mid1 = 12MO head, mid2 = 18MO head, aged = 32MO head, amp1/amp2/amp3 = heads regenerated from 32MO animals.

**Supplementary Table 4 Cell type annotation with published markers**

Table4.1 Sub-clusters of neoblast, neurons, muscles, and cathepsin+ cells were assigned to cell types according to specific expressions of certain markers.

Table4.2 References from which the cell type markers were collected.

**Supplementary Table 5 Cell proportions and Cell cycle proportions**

Table5.1 Cell counts and proportions of neoblast cell types in each sample at different cell cycle phases.

Table5.2-9 Cell counts and proportions of neoblasts, neurons, muscle, protonephridia, cathepsin+ cells, epidermis, intestine, and secretory cell types in each sample and at different cell cycle phases.

Table5.10 Cell counts and proportions of each tissue type in each sample.

**Supplementary Table 6 es-neo stem cell cell cycle proportions**

Cell proportions of sub-cluster of es-neo at different cell cycle phases.

**Supplementary Table 7 Cell clusters with high vulnerability to age-associated transcriptional changes**

Summary information of vulnerable sub-clusters and control sub-clusters, including DEG number, percentage of piwi1+ cells, mean CytoTRACE score, abundance in the tissue, and cell proportion at each cell cycle phase in different conditions.

**Supplementary Table 9 Cell cycle markers used in cell cycle analysis**

Gene markers of S phase and G2M phase with planarian homologs.

**Supplementary Figure 1** Characterization of three neoblast subtypes, sigma, zeta, and gamma, based on their cluster-enriched expression of soxp-1, soxp-2, soxp-3, zfp-1, egr-1, p53, hnf4, gata456-1, and nkx2.2.

**Supplementary Figure 2** Characterization of the epidermal lineage. **a**, UMAP plot of epidermis sub-clusters and zeta neoblasts. **b**, Cluster-enriched gene expression of epidermal lineage markers, zeta (zfp-1) and early epidermal progenitors (p53 and prog-1). **c**, UMAP plot colored by cell cycle phase of epidermal lineage cells.

**Supplementary Figure 3** Characterization of the intestine lineage. **a**, UMAP plot of intestine sub-clusters and gamma neoblasts. **b**, Cluster-enriched gene expression of intestine markers (apob-1 and apob-2), proliferating markers (mcm4, mcm5, and mcm7), and post meiotic markers (xbp1 and p4hb). **c**, UMAP plot colored by cell cycle phase of intestine cells.

**Supplementary Figure 4** Gene expression of tgs-1 and tspan-1 in neoblasts.

**Supplementary Figure 5** foxF-1 expression pattern in the neoblast sub-populations.

**Supplementary Figure 6** Violin plots showing expression level of the muscle markers in muscle cell types.

**Supplementary Figure 7** Cluster-enriched gene expression of muscle differentiation-related TFs in neoblast population.

**Supplementary Figure 8** Specific expression of cathepsin+ cell markers in neoblasts (**a**) and cathepsin+ cells (**b**).

**Supplementary Figure 9** Specific expression of F-spondin, LDLRR-3, and ETS-1 in phi and beta neoblasts.

**Supplementary Figure 10** Violin plots showing differential expression level of foxF-1, nf1c, ptprj, F-spondin, and LDLRR-3 between phi and beta neoblast.

**Supplementary Figure 11** Restricted expression of yolk gland-related genes in CPEB1+ neoblasts.

**Supplementary Figure 12** Violin plots exhibiting extensive expression of different differentiated lineage markers in es-neo.

**Supplementary Figure 13** Cluster-enriched expression of epidermal markers in epidermal lineage cells with zeta included.

